# FoldMapper: Protein Inverse Folding Model with Structure-Derived Evolutionary Profiles and a Multiscale Graph Encoder

**DOI:** 10.64898/2026.11.15.766879

**Authors:** Haelyn Kim, Juyong Lee

## Abstract

Protein inverse folding designs amino-acid sequences that fold into a target backbone structure. Existing deep-learning methods exploit backbone geometry but not evolutionary statistics. Those statistics come from a multiple sequence alignment, which must be built from a query sequence. That query is the sequence to be designed, so the dependency is circular. We present FoldMapper, an inverse folding model that breaks this circularity. It retrieves structural homologues by shape alone, then aggregates their amino-acid identities into a per-residue structural position-specific frequency (SPSF) profile. A Multiscale Graph Encoder passes a sparse profile-conditioned encoding into a dense structure-conditioned message-passing network. FoldMapper reaches 60.0% best-of-100 sequence recovery on TED domains, 64.9% on PDB, and 51.3%/53.1% on PPIBindBench non-peptide and peptide binders. Across both benchmarks, it lies on the recovery–diversity Pareto front. Structure self-consistency surpasses all baselines on RFdiffusion de novo monomer and binder backbones, and matches the strongest baselines on natural proteins. Ablations and profile-quality analyses show that structure itself encodes recoverable evolutionary information, accessible without a sequence-homology search.

## 1 Introduction

Protein inverse folding models aim to design sequences that fold into a target backbone structure [1–3], which is a central task in computational protein design [4, 5]. A general protein design pipeline runs in three steps. It generates protein-like backbones, designs sequences expected to fold into them, and validates the designs in silico by comparing designed and predicted structures [6, 7]. Recent deep learning approaches formulate inverse folding as a conditional sequence generation task over given protein backbone structures. Ingraham et al. [8] first proposed the geometric graph-based sequence generative model for protein design. ProteinMPNN [4] extended this approach with the message-passing neural network (MPNN) architecture [9] and autoregressive decoding. Subsequent sequence generative methods incorporated additional information, such as protein language model representations [5, 10], knowledge-refined features [11], and diffusion-based generative frameworks [12], achieving higher sequence recovery rates. These methods are widely adopted as essential tools for de novo protein design.

Existing protein sequence generation methods rely solely on geometric features extracted from an input backbone structure while ignoring evolutionary information. Multiple sequence alignments capture co-evolutionary couplings between residues, which encode spatial information [13–15]. Early work used mutual information to predict residue contacts [16]. Later work advanced to statistical coupling analysis and direct coupling analysis [17, 18]. However, the earlier methods suffered from high false-positive rates. Eventually, AlphaFold and other recent structure prediction models proved that deep-learning algorithms capture and exploit this information successfully leading to near-experimental accuracy in structure prediction [19–21].

We hypothesised that the identical principle applies in the inverse folding problem. If evolutionary information is critical for predicting structure from homologous sequences, generating sequences for a given backbone structure would benefit from sequence information of homologous structures. Evolutionary information is normally read from a multiple sequence alignment (MSA), which is built by searching with the query sequence [22, 23]. In inverse folding that query is the designed sequence itself, so the search is circular. In this study, we propose FoldMapper, an inverse folding model that integrates evolutionary information from structurally homologous proteins via a Multiscale Graph Encoder (MGE). The MGE feeds a profile-conditioned graph encoding into a structure-conditioned MPNN. Structural homologs are retrieved with Foldseek [24], which searches on shape alone, without sequence information. Their pairwise alignments are stacked on the query to give a per-residue amino-acid frequency, an approximation of multiple structure alignment [25–27]. The amino acid identities at each aligned position across these homologs form a per-residue structural position-specific frequency (SPSF) profile. This profile is the structure-derived analogue of a sequence PSSM [22]. It encodes the evolutionary sequence preferences of the backbone fold. The SPSF profile is thus an evolutionary profile derived from structure rather than from a sequence search.

For rigorous benchmark of FoldMapper, we compiled two new benchmark datasets: one for single-domain and the other for multimeric backbone structures. For single-domain backbone structures, we assembled TEDMonoBench by augmenting experimentally resolved structures with The Encyclopedia of Domains (TED) domains [28] curated from the AlphaFold database [29]. Sequence generation of multimeric backbone structures is becoming more important for the development of biotherapeutics, such as mini-binders and antibodies. To evaluate sequence generation performance on heterodimeric backbone structures, we compiled the PPIBindBench dataset. PPIBind-Bench is a leakage-controlled benchmark of 576 non-redundant heterodimer complexes from recent PDB entries, spanning peptide and non-peptide binders held below 30% sequence identity to the training set. Across both benchmarks, FoldMapper achieves high sequence recovery while maintaining comparable sequence diversity and structure self-consistency. It attains the highest best-of-100 sequence recovery on TED (60.0%) and peptide binders (53.1%), and the second highest on PDB (64.9%) and non-peptide binders (51.3%). It also lies on the recovery–diversity Pareto front. These results show that incorporating evolutionary information from SPSF profiles improves the accuracy of protein inverse folding.

## 2 Results

### 2.1 The FoldMapper architecture

FoldMapper is a multiscale graph neural network for protein inverse folding that integrates SPSF profiles into a message-passing framework [4, 9] (Fig. 1). The Multi-scale Graph Encoder (MGE) constructs two independent *k*-nearest neighbour (*k*-NN) graphs over the residue C*α* atoms of the same backbone. Passing the profile directly to the structure decoder provides a limited signal, as the ablation confirms (Section 2.6). The MGE therefore adds a dedicated encoder that first processes the profile in local backbone context, producing stronger node features for decoding. A profile-conditioned encoder (Stage 1) first produces evolutionary-aware node embeddings. These embeddings initialize a structure-conditioned MPNN (Stage 2) that decodes the sequence autoregressively. The two graphs differ in neighbourhood size. Stage 1 uses a sparser local graph (*k*=30), and Stage 2 a denser graph (*k*=48) that reaches longer-range contacts. The model accepts the backbone atomic coordinates (N, C*α*, C, O) at each residue position *i* as input.

**Fig. 1:**
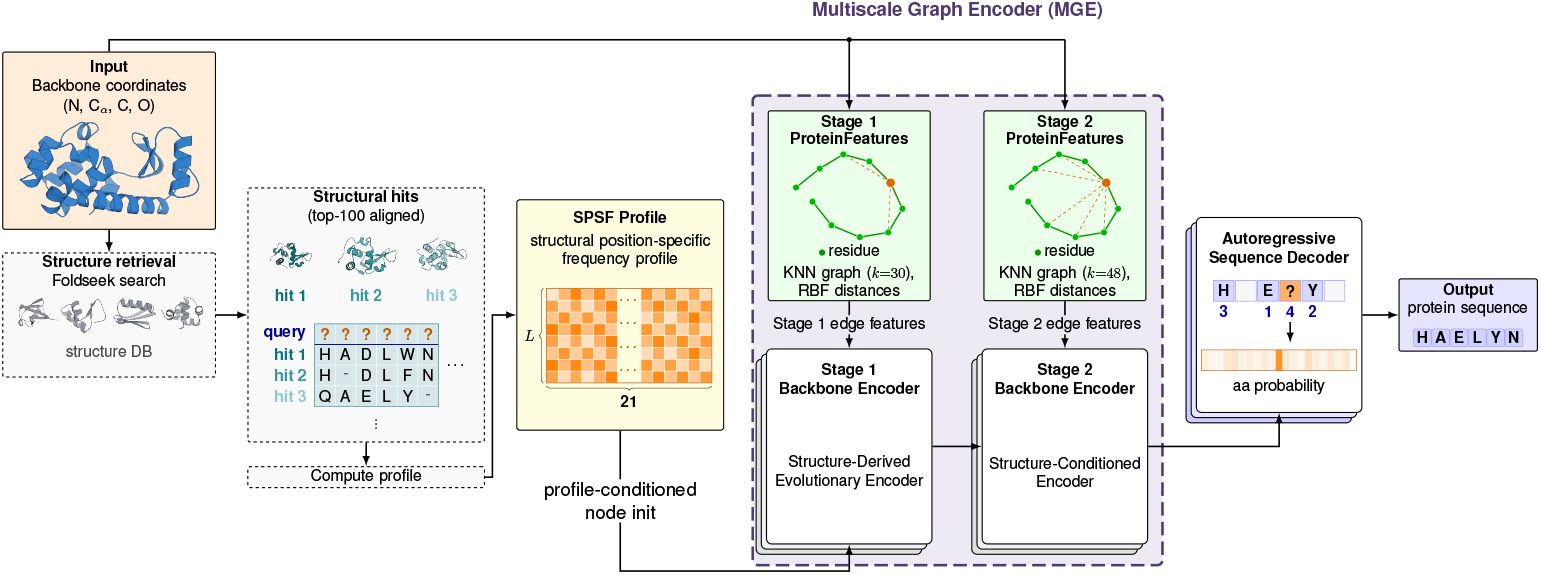
FoldMapper architecture. A backbone structure enters at the left and leaves as a designed sequence at the right. Foldseek retrieves structural homologues from a structure database, and the top 100 aligned hits are accumulated into the *L* × 21 structural position-specific frequency (SPSF) profile, shown as a heatmap. The purple dashed outline encloses the Multiscale Graph Encoder (MGE), which builds two *k*-nearest-neighbour graphs over the same backbone: *k*=30 for Stage 1 and *k*=48 for Stage 2. Each stage derives its own edge features from radial basis functions over interatomic distances, and feeds a stack of message-passing layers drawn as stacked boxes. The profile enters only at Stage 1, as the profile-conditioned node initialisation. Stage 2 conditions on backbone geometry alone. Its node representations pass to the autoregressive decoder, which emits one amino-acid distribution per position under a random decoding order, numbered above the residues. Hidden dimensions, layer counts and training details are in Methods.

### 2.2 Sequence recovery and diversity

Benchmark results show that FoldMapper achieved the highest or second highest sequence recovery for both monomeric and heterodimeric structures while maintaining moderate sequence diversity (Table 1). For the high-quality predicted backbone structures (TED), FoldMapper models achieved the highest sequence recovery of 60.0%,

**Table 1:** Recovery, perplexity, and diversity across benchmarks.

| TEDMonoBench | **TED** (*n*=10,000) |  |  | **PDB** (*n*=2,126) |  |  |
|---|---|---|---|---|---|---|
|  | Rec. | Ppl. | Div. | Rec. | Ppl. | Div. |
| ProteinMPNN | 46.6 | 5.93 | 28.9 | 50.3 | 5.14 | 28.0 |
| ESM-IF | 56.3 | 4.07 | 12.6 | 59.3 | 5.61 | 11.5 |
| KWDesign | 55.7 | 4.61 | 5.1 | **65.2** | 19.55 | 3.7 |
| MapDiff (*n*=10,000/2,119) | 43.2 | 13.25 | **54.6** | 42.5 | 14.15 | **52.3** |
| LM-Design (*n*=10,000/2,125) | 51.6 | 5.25 | 16.1 | 64.2 | 5.35 | 11.3 |
| FoldMapper-29.5M | **60.0** | 3.79 | 20.1 | 64.9 | **3.63** | 13.8 |
| FoldMapper-40.0M | **60.0** | **3.78** | 19.1 | 63.0 | 3.79 | 15.4 |
| FoldMapper-29.5M+BLOSUM | 52.2 | 3.80 | 25.2 | 53.5 | 3.72 | 25.4 |
| FoldMapper-40.0M+BLOSUM | 53.7 | 3.80 | 21.4 | 58.4 | 3.70 | 17.9 |
| PPIBindBench | **Non-peptide** (*n*=197) |  |  | **Peptide** (*n*=379) |  |  |
|  | Rec. | Ppl. | Div. | Rec. | Ppl. | Div. |
| ProteinMPNN | 46.2 | 6.38 | 26.7 | 51.0 | 8.42 | 20.0 |
| ESM-IF (*n*=141/369) | 51.0 | 5.37 | 14.8 | 41.9 | 11.17 | 15.1 |
| KWDesign | **54.4** | **4.76** | 6.6 | 52.5 | **7.81** | 8.0 |
| MapDiff (*n*=191/377) | 34.3 | 26.2 | **60.6** | 38.5 | 71.9 | **74.4** |
| LM-Design (*n*=196/379) | 31.7 | 5.51 | 16.4 | 21.0 | 17.71 | 11.4 |
| FoldMapper-29.5M | 51.3 | 5.73 | 21.2 | **53.1** | 8.41 | 17.4 |
| FoldMapper-40.0M | 50.3 | 5.83 | 21.1 | 52.2 | 8.36 | 17.2 |
| FoldMapper-29.5M+BLOSUM | 45.5 | 5.61 | 28.0 | 50.1 | 8.32 | 21.9 |
| FoldMapper-40.0M+BLOSUM | 48.3 | 5.73 | 24.4 | 51.6 | 8.28 | 20.9 |

Recovery is best-of-100 per structure and diversity is the mean over all 100 samples. Perplexity is teacher-forced on the native sequence, so it does not depend on sampling (Methods). Each metric is then averaged over proteins (*T*=0.1).
Best results in **bold**; second-best underlined.
Supplementary Table S1 reports mean-of-100 recovery; Supplementary Table S2 keeps best-of-100 but takes the median across proteins.

whereas other methods yielded recovery ranging from 43.2 to 56.3%. The diversity of generated sequences with FoldMapper showed 20.1%, which is higher than that of ESM-IF and LM-Design, 12.6% and 16.1%, respectively. This places FoldMapper on the recovery–diversity Pareto front, outperforming baseline methods that suffer from a steep trade-off between the two metrics (Fig. 2a). Together with FoldMapper, Protein-MPNN and MapDiff also form the recovery–diversity Pareto front with lower recovery and higher diversity. On the monomeric crystal structures, the PDB set, FoldMapper placed a close second with recovery 64.9%, 0.3 percentage points below KWDesign. However, FoldMapper showed much higher diversity of 13.8% compared to 3.7% of KWDesign. These results demonstrate that FoldMapper generates highly-probable sequences while searching sequence space widely.

**Fig. 2:**
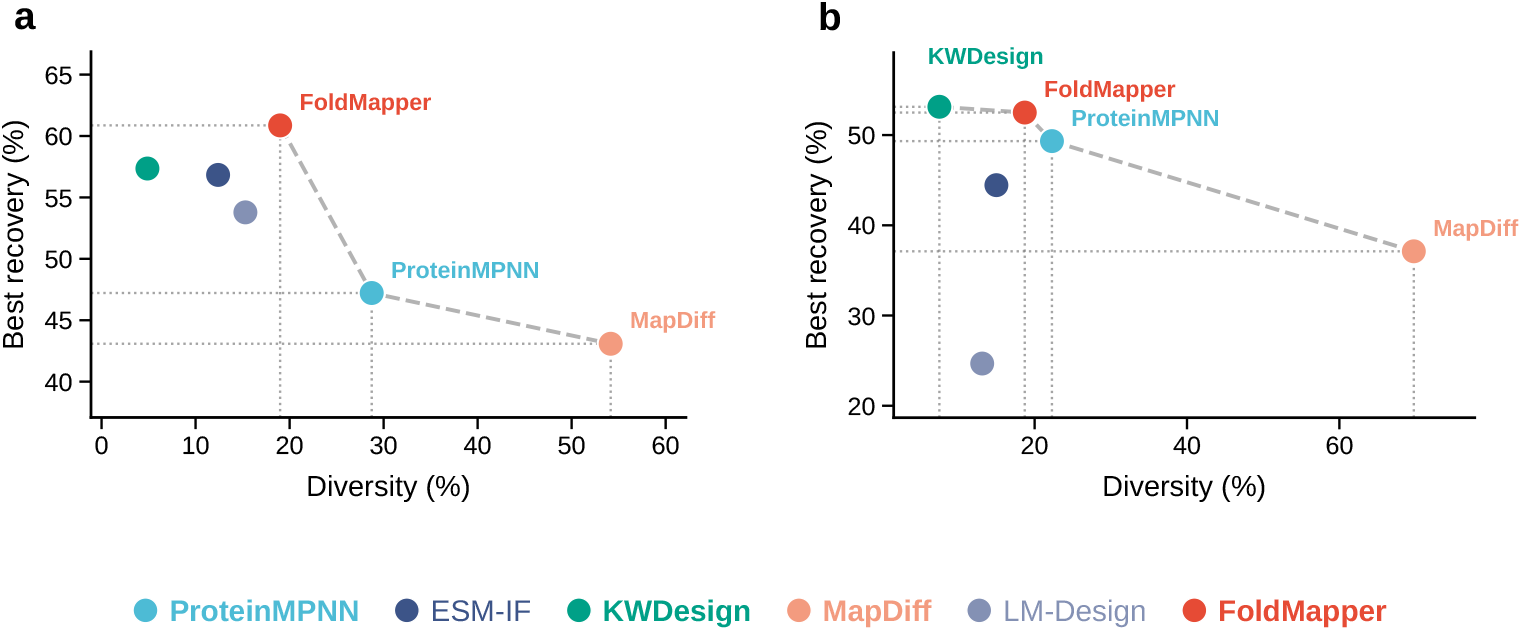
Recovery–diversity comparison across benchmarks. Best-of-100 sequence recovery versus mean pairwise diversity (*T* =0.1, 100 samples per structure), pooled over both subsets of each benchmark. FoldMapper uses SPSF profiles with every 100%-identity hit to the query excluded (Methods). **a**, TEDMonoBench (TED, *n*=10,000; PDB, *n*=2,126). **b**, PPIBindBench heterodimer complexes filtered at ≤30% sequence identity to training data (Non-peptide, *n*=197; Peptide, *n*=379). Dotted guides drop from each Pareto-optimal point to the axes. The dashed line connects the Pareto front (FoldMapper, ProteinMPNN and MapDiff in **a**; these three and KWDesign in **b**).

For the binder sequence design benchmark, PPIBindBench, FoldMapper also yielded top-tier sequence recovery with reasonable diversity (Fig. 2b). FoldMapper achieved the highest sequence recovery for peptide binders, 53.1%, and the second highest for non-peptide binders, 51.3%. For both binder design tasks, KWDesign [11] reached competitive recovery but resulted in the lowest sequence diversity among all tested methods, at most 8.0% on every subset, which limits efficient sequence-space exploration and protein design. FoldMapper produced more diverse sequences than ESM-IF and LM-Design on every benchmark subset. On non-peptide binders, ESM-IF was performed on only the 141 of 197 entries due to memory limitation. For those 141 entries, FoldMapper still showed higher recovery of 52.8% than that of ESM-IF, 51.0%. These results show that incorporating evolutionary information carried by SPSF profiles improves the accuracy of sequence generation for given backbone structures.

The results show that increase in parameter size, the hidden dimension of the encoder, has little effect on sequence generation performance. Among the FoldMapper variants, the FoldMapper-29.5M model without the BLOSUM62 prior attains the highest recovery on PDB and both PPIBindBench subsets (Table 1). On TED, it matches the 40.0 M model at 60.0%. This indicates the SPSF profile already conveys sufficient substitution information. These results indicate that the MGE with SPSF profiles yields a favourable recovery–diversity balance that no single tested baseline achieves at comparable recovery.

### 2.3 Structure self-consistency benchmark on de novo backbones

De novo backbones have no native sequence to recover, making them a stringent, design-relevant test of whether a model specifies a foldable sequence for a novel structure. We evaluated each model on the 758 RFdiffusion generated de novo backbones (Section 4.5). Designs were folded with two different folding methods: AlphaFold 3 [21] and Boltz2 [30], which yielded consistent rankings (Supplementary Table S8).

The leading FoldMapper models surpass all baselines in generating high-quality foldable sequences of monomer and binder backbones (Fig. 3a and b). On the monomer backbones, FoldMapper-generated sequences show higher success rates than the tested baselines (Fig. 3a, left). FoldMapper-29.5M and 40.0M achieved notably high success rates of 91% and 95%, respectively, indicating that most generated sequences folded to the input structure. Among the tested baselines, ESM-IF showed the best success rate of 59% followed by that of KWDesign, 44%.

**Fig. 3:**
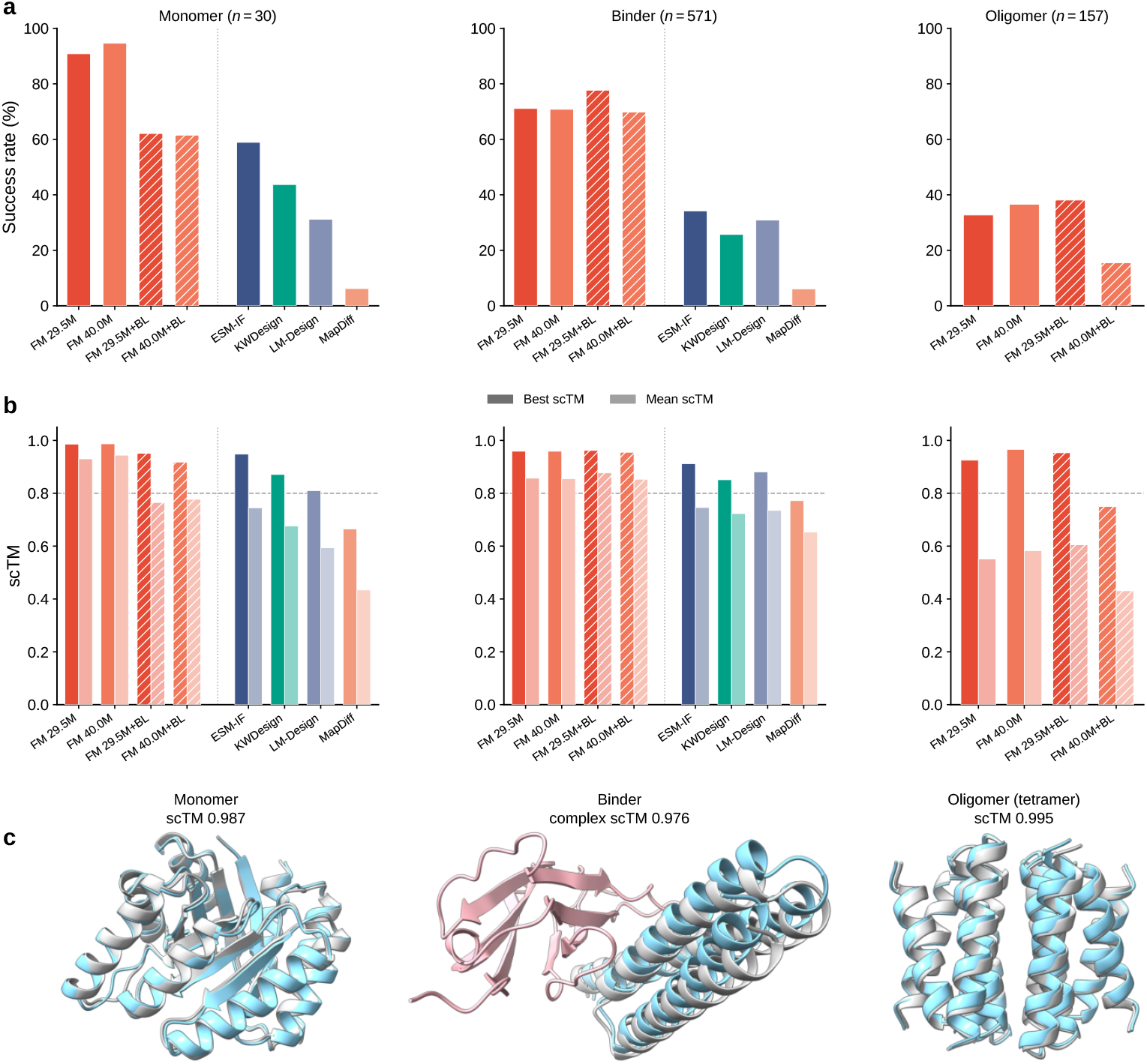
Structure self-consistency and representative designs on RFdiffusion de novo backbones. **a**, Success rate on RFdiffusion de novo backbones (AlphaFold 3 [21]), the fraction of 100 samples with scTM ≥ 0.8. **b**, Best-of-100 scTM (dark bars) and the per-entry mean (light bars). The dashed line marks the scTM = 0.8 threshold that defines the success rate in **a**. Columns are the three subsets throughout: monomer (*n*=30), binder (*n*=571) and oligomer (*n*=157). scTM is complex scTM for the multi-chain binder and oligomer targets. Bars are means over entries and carry no error bars. The dotted vertical line separates FoldMapper variants from baselines. FoldMapper bars are labelled by trainable parameter count: dark orange = 29.5 M, light orange = 40.0 M. Solid = without BLOSUM, hatched = +BLOSUM. The oligomer column is restricted to models supporting tied sequence design. ProteinMPNN is excluded because the benchmark backbones were screened with ProteinMPNN sequences, and is reported in Supplementary Table S8. **c**, Representative best-of-100 designs (RFdiffusion backbone in grey, FoldMapper prediction in cyan, fixed target chains in pink): monomer (`TIM_exptested_design_3`, 227 aa), scTM 0.987; TrkA binder (`Trka_binder_design46`), complex scTM 0.976; symmetric tetramer (`HE0427`), scTM 0.995.

The FoldMapper models also outperform the other methods in terms of achieving high self-consistency between the input and folded structures, a metric for quality of generated sequences (Fig. 3b). FoldMapper-40.0M leads on best-of-100 scTM with 0.987, followed by FoldMapper-29.5M with 0.986. The best performing baseline on the monomer backbones is ESM-IF with a best scTM of 0.948. The other methods, KWDesign, LM-Design and MapDiff show much lower self-consistency with best-of-100 scTM values of 0.871, 0.810, and 0.665, respectively. Larger gaps are observed in terms of mean scTM than best-of-100. FoldMapper-29.5M keeps a high mean scTM, 0.930 on monomers, whereas the other baselines show much lower mean scTM due to low success rates. ESM-IF, the best baseline, shows the mean scTM of 0.745.

On binder complex backbones, all FoldMapper models surpassed the baselines by a wide margin (Fig. 3a and b, middle). FoldMapper-29.5M reached a success rate of 71% with a best scTM of 0.959. All four FoldMapper models exceeded a best scTM of 0.95 and mean scTM of 0.8. Contrary to the monomer benchmark, adding the BLOSUM prior led to a small performance gain on binders for the 29.5 M model. FoldMapper-29.5M+BLOSUM reached the highest success rate of 78% among all tested models. The success rates of the other baselines remained below 40%. The strongest baseline, ESM-IF, resulted in a success rate of 34% and a best scTM of 0.912.

Generating sequences for symmetric oligomer backbones is a more challenging task because tied sequence generation is required. The three leading FoldMapper models reached 33–38% success and 0.925–0.966 best scTM, while FoldMapper-40.0M+BLOSUM lagged (Fig. 3a–c, right). The mean-to-best gap reaches 0.374 even for FoldMapper (0.551 against 0.925). Thus, rare foldable samples dominate the result (Fig. 3b). Note that the other baseline methods were excluded from the analysis because they do not support the tied generation.

The high accuracy of FoldMapper models on monomer, binder and oligomer back-bones indicates that evolutionary information derived from the SPSF profile facilitates generating foldable sequences for de novo structures. Representative cases across the three different tasks are illustrated in Fig. 3c. In each case, the AlphaFold 3 prediction superimposes on the input backbone at scTM 0.976 or above, demonstrating high-quality of FoldMapper generated sequences.

### 2.4 Structure self-consistency on native-like backbones

We next investigated whether the high accuracy of FoldMapper retains for native and native-like backbone structures. We evaluated the sequence generation results on 101 TEDMonoBench structures and 100 PPIBindBench complexes (Supplementary Methods). The benchmark results show that all baselines perform better than on de novo generated backbones and the FoldMapper models maintain top-tier performance on native-like backbones. For the monomer structures, the FoldMapper models achieved the second-best success rates on the TED and PDB sets, closely trailing ProteinMPNN and ESM-IF, respectively. In terms of complex structures, FoldMapper-29.5M+BLOSUM achieved the highest success rate for non-peptide binders under AlphaFold 3 and the second highest under Boltz2. Peptide binders showed little separation among models under either predictor, spanning 91.5–95.0% and 94.6–96.0%. These results demonstrate that FoldMapper generates high-quality sequences on native-like as well as de novo backbones. Full per-subset metrics and success rates are in Supplementary Tables S6 and S7, and PAE_int_ in Supplementary Table S7.

We extended the same evaluation to 30 antibody–antigen complexes, jointly designing the heavy and light chains. FoldMapper performs comparably to the antibody-specific model AbMPNN [31] under AlphaFold 3 but trails it under Boltz2. It outperforms AntiFold [32] on complex scTM, success rate and DockQ under both predictors (Supplementary Table S9). However, success rates remain below 30% and mean DockQ [33] values below 0.15 for all methods. These results suggest that SPSF profiles support competitive antibody sequence design without antibody-specific training, although reliable interface recovery remains challenging.

### 2.5 SPSF profile quality analysis

To understand how SPSF profile is used to improve FoldMapper’s performance, we analyzed the relationship between three profile characteristics (hit quality, profile sharpness, and hit count) and per-protein sequence recovery. More detailed metric definitions are described in Supplementary Methods.

It is demonstrated that the quality of the best homologous hit is critical in high sequence recovery (Fig. 4). The bit score of the best Foldseek hit, the quality of aligned structures, correlates most with average sequence recovery with a Spearman *ρ*=+0.281. Recovery in this subsection is the mean over the 100 samples per structure, not the best-of-100 reported elsewhere. Stratifying proteins by max bit score quintile reveals a monotonic increase from 53% recovery in the lowest quintile to 63% in the highest (Fig. 4a). Generally, sharper profiles lead to higher sequence recovery (Fig. 4b). Lower Shannon entropy correlates with higher recovery, though less strongly than the bit score (*ρ*=− 0.152), with the sharpest entropy quintile reaching 61% versus 54% for the flattest. Both correlations persist after controlling for protein length, Spearman *ρ*= + 0.185 and *ρ*= − 0.115, respectively (Supplementary Table S4). In contrast, the number of homologous hits contributes little to sequence recovery, demonstrated by a Spearman *ρ*= + 0.034 (Fig. 4c). Proteins with no hits show much lower recovery (25%), but recovery plateaus at 53–56% once at least one hit is retrieved. The collapse to 25% recovery without any hit shows that homolog search plays a critical role in the performance of FoldMapper.

**Fig. 4:**
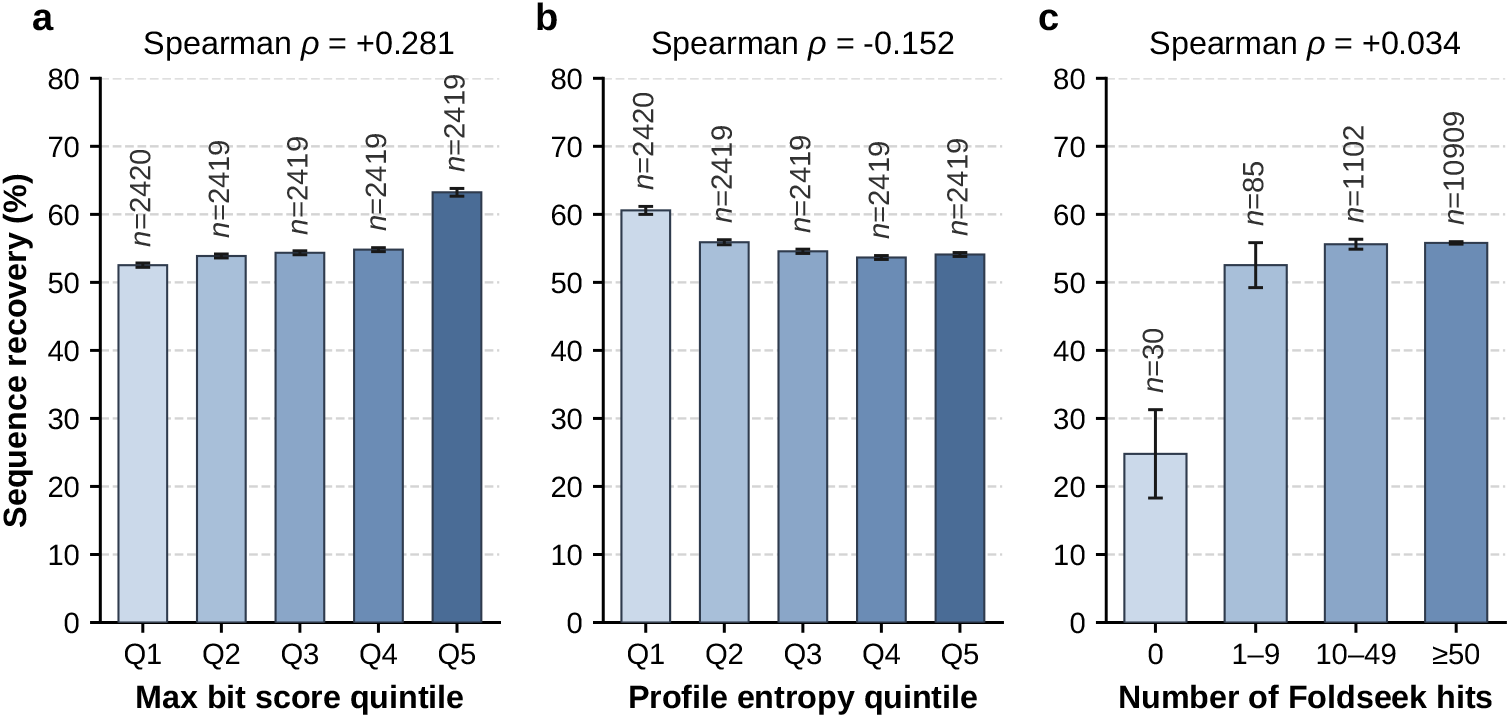
The relationship between characteristics of the SPSF profile and sequence recovery. Sequence recovery on TEDMonoBench (TED and PDB pooled) stratified by (**a**) max Foldseek bit score quintile, (**b**) mean Shannon entropy quintile of the SPSF profile, and (**c**) number of Foldseek hits. Recovery is the mean over the 100 sequences sampled per structure, and quintiles hold equal numbers of proteins. Panels a and b need both a bit score and an entropy, so they use *n*=12,096 proteins. Panel c uses the full benchmark (*n*=12,126), because proteins without any hit have neither metric. Error bars denote 95% confidence intervals, and per-bar *n* is given above each bar. Hit count bins: 0, 1–9, 10–49, ≥50. Profiles exclude every hit at 100% sequence identity to the query, as everywhere else in this work (Methods).

Overall, our analysis indicates that the quality of the best homolog is the strongest correlate of high sequence recovery, not the depth of the profile. This contrasts with the protein folding problem. In structure prediction, a deeper MSA generally provides stronger co-evolutionary signal and higher accuracy [21]. However, in the inverse folding problem, the existence of a structural homolog is essential rather than the number of retrieved homologs. This can be attributed to the different nature of folding and inverse folding problems. Sequence information is high entropy and high degeneracy with respect to a folded structure. Protein folding compresses vast sequence space into more constrained structure space. Thus, for structure to sequence mapping, small variations in structure may lead to larger shift in sequence space. This suggests that interpreting or extracting information from multiple structure alignment may require greater care.

### 2.6 Ablation analysis

To identify the origin of FoldMapper’s performance gains, we performed ablation analysis of two core design choices: (i) the SPSF profile input, and (ii) the Stage 1 encoder of the Multiscale Graph Encoder (MGE), whose removal leaves the single Stage 2 encoder (Table 2 and Supplementary Table S3). In the single-stage variant, the profile-conditioned node initialisation serves as the Stage 2 initial node features, without a separate Stage 1 encoder. The “w/o Stage 1 encoder & profile” variant removes both, training the Stage 2 MPNN layers from scratch on the same training set.

**Table 2:** Ablation of the SPSF profile and the Stage 1 encoder.

| TEDMonoBench | **TED** (*n*=10,000) |  |  | **PDB** (*n*=2,126) |  |  |
|---|---|---|---|---|---|---|
| Variant | Rec. | Ppl. | Div. | Rec. | Ppl. | Div. |
| FoldMapper | **60.0** | **3.79** | 20.1 | **64.9** | **3.63** | 13.8 |
| w/o SPSF profile | 59.1 | 3.88 | 21.5 | 52.2 | 4.90 | 24.0 |
| w/o Stage 1 encoder | 53.8 | 4.60 | 25.1 | 63.4 | 3.77 | 18.9 |
| w/o Stage 1 encoder & profile | 52.8 | 4.76 | 26.0 | 50.2 | 5.16 | 27.1 |
| PPIBindBench | **Non-peptide** (*n*=197) |  |  | **Peptide** (*n*=379) |  |  |
| Variant | Rec. | Ppl. | Div. | Rec. | Ppl. | Div. |
| FoldMapper | **51.3** | **5.73** | 21.2 | **53.1** | **8.41** | 17.4 |
| w/o SPSF profile | 46.9 | 6.24 | 24.3 | 52.6 | 8.67 | 17.4 |
| w/o Stage 1 encoder | 50.5 | 5.85 | 23.9 | 52.0 | 8.53 | 19.5 |
| w/o Stage 1 encoder & profile | 45.8 | 6.46 | 26.8 | 52.0 | 8.97 | 19.0 |

Sequence recovery, perplexity, and diversity when removing each core component. Aggregation follows Table 1 (*T*=0.1, 100 samples per structure).
The FoldMapper and “w/o Stage 1 encoder” rows use SPSF profiles with every 100%-identity hit excluded (Methods). The two profile-free rows are unaffected by that filter.

Removing the SPSF profile reduces the sequence recovery of the predicted structures by 0.9 pp from 60.0% → 59.1%. Interestingly, the decrease is far larger on the PDB set (from 64.9% → 52.2%, 12.7 pp). Without the profile, sequence recovery on the TED set even surpasses that on the PDB set. Thus, the profile’s benefit concentrates on the PDB set, where strong structural homologs are available (Supplementary Table S5).

The results show that the Stage 1 encoder affects most on the TED set and marginally on the PDB set. Removing the Stage 1 encoder reduces sequence recovery on the TED set by 6.2 pp, from 60.0% to 53.8%. On the PDB set the drop is 1.5 pp, from 64.9% to 63.4%. This difference implies that the proposed MGE architecture may be critical in refining the geometric distribution shift and the weaker profile information of predicted domains via hierarchical information exchange. Stage 1 is the only module that reads the SPSF profile, and it exchanges the profile information across each residue’s local backbone neighbourhood (*k*=30). Stage 2 adds no further profile input and refines the Stage 1 node features over a wider backbone graph (*k*=48). Thus, the single-stage variant tests the effect of local mixing, not the profile itself. Removing both components yields sequence recovery of 52.8% on the TED set and 50.2% on the PDB set, corresponding to decreases of 7.2 and 14.8 pp from FoldMapper.

On the non-peptide binders of PPIBindBench, profile removal led to a notable decrease in sequence recovery of 4.4 pp, which also shows the importance of structure alignment information. Removing the Stage 1 encoder as well costs 5.5 pp, the largest drop on this subset. On the contrary, peptide binders show only marginal changes (≤2 pp) across all ablations. These results indicate that the profile appears to add little information for these short binders (≤30 aa). Recovery remains near 52–53% and derives mainly from the interface geometry rather than from evolutionary signal. Short fragments rarely retrieve informative structural homologs, leaving little evolutionary signal to remove. Chain length and homolog strength are coupled within TEDMonoBench as well. TED chains of 50 aa or fewer retrieve a best hit scoring 76 on average, against 359 for chains longer than 250 aa (Supplementary Table S5).

## 3 Discussion

We developed a new protein inverse folding model FoldMapper that bridges two paradigms of existing models: structure-only autoregressive models [4, 8] and sequence-or MSA-based language models [5, 10, 11]. FoldMapper contributes two elements: the SPSF profile derived from structural homologs, and the two-stage MGE architecture. In contrast to most existing inverse folding models that consider only backbone geometric information, FoldMapper incorporates the sequence–structure relationship carried by SPSF profiles. This frees inverse folding from regimes where sequence homologues are scarce or absent, as demonstrated on de novo backbones [6]. Potential applications extend to orphan families and engineered scaffolds [34].

The benchmark results demonstrate that FoldMapper outperforms most tested baselines on sequence recovery while maintaining the diversity of generated sequences. KWDesign is the only baseline to exceed FoldMapper’s recovery, on PDB by a slight margin and on non-peptide binders. It reaches that recovery by refining its design with two frozen pretrained models, ESM2-650M and ESM-IF, over three recycling passes [11]. That recovery costs diversity: FoldMapper’s is three to four times higher on both subsets. Aggregated over TEDMonoBench, FoldMapper leads on both axes and KWDesign leaves the recovery–diversity front (Fig. 2a). On PPIBindBench neither model dominates the other, and both lie on it (Fig. 2b). The same balance pays off on de novo backbones: FoldMapper surpasses all baselines on monomer and binder backbones. On natural proteins, FoldMapper remains competitive with the strongest baselines. On antibody sequence design tasks, the quality of FoldMapper generated sequences matches AbMPNN under AlphaFold 3 and trails it under Boltz2, without antibody-specific training.

The analysis of SPSF profile characteristics on TEDMonoBench shows that higher similarity of the nearest homolog and lower profile entropy are associated with higher sequence recovery. Also, the existence of a structural homolog is more important than the number of homologs. In summary, FoldMapper’s performance depends on the presence and quality of the nearest structural homolog rather than profile depth. Ablation analysis confirms that both the SPSF profile and the multiscale encoder contribute. The profile matters most on crystal structures and the encoder most on predicted domains. Removing either component reduces recovery, and removing both reduces it further except on peptide binders. Together, these analyses suggest that SPSF profiles carry rich evolutionary information and can be a foundation of next-generation sequence design or co-design models.

Two limitations follow from the architecture. First, FoldMapper depends on retrieving a structural homolog, and recovery falls to 25% when Foldseek returns no hit (Section 2.5). The profile is then the zero vector, which is the condition the model is trained for. Therefore, it falls back to structure-only prediction. Second, FoldMapper requires a structure homology search before inference (Methods).

One of the directions for further improvement is to use domain-specific homology search, i.e., searching against only Ig folds for antibody design. Using such a focused database could supply cleaner evolutionary signal for CDR sequence selection without antibody-specific training. The same principle could be extended to enzyme scaffolds, membrane-protein topologies, and other specialised design tasks.

## 4 Methods

### 4.1 Dataset preparation

To improve the accuracy and generalizability of sequence generation, we augmented the ProteinMPNN dataset [4] with interacting domain pairs from TED [28]. A TED interacting superfamily pair (ISP) couples two domains that pack together within the same AFDB chain [29]. TED assigns each domain a consensus confidence from the agreement of its three domain-parsing methods. High means all three agree on the boundary, Medium that two agree. We kept pairs with inter-domain PAE ≤ 4 and Medium or High consensus confidence, yielding 32,271,842 unique domains after deduplication. These were clustered at 50% sequence identity using MMseqs2 [35]. A single representative per cluster was selected, giving 1,547,541 domains.

This filtered TED dataset was merged with the ProteinMPNN dataset consisting only of crystal structures. The combined set was clustered at 50% sequence identity using MMseqs2 to define homology-level partitions, resulting in 1,582,548 clusters. We assigned every cluster containing a ProteinMPNN test chain to the primary test split. These 2,114 test clusters hold 18,573 chains, from which 2,126 were retained as the PDB evaluation set. The remaining 1,580,434 clusters were split 9:1 into 1,422,390 training and 158,044 validation clusters. This cluster-level split guarantees no homologous leakage.

To test generalization more rigorously, we assembled the second evaluation set comprising 10,000 TED domains. These domains were drawn from the held-out range 4 *<* PAE ≤ 5, disjoint from the PAE ≤ 4 training domains. At Medium or High consensus confidence this range yielded 11,651,363 domains. A first-pass sequence search removed domains matching the training, validation, or primary test sequences. The remaining domains were clustered at 30% identity and 70% bidirectional coverage, leaving 260,278 representatives. The representatives were then clustered again at the same thresholds, together with all training, validation, and primary test sequences. Any representative sharing a cluster with a reference sequence was discarded, leaving 162,417 candidates. From this pool, 10,000 domains were drawn by length-stratified sampling into 20 bins of 500, at most one domain per cluster. The released benchmark lists every identifier. The final evaluation set consisted of 2,126 PDB chains from the held-out test clusters combined with the 10,000 length-stratified TED domains. We refer to this combined monomeric benchmark as TEDMonoBench.

### 4.2 PPIBindBench benchmark construction

To evaluate inverse folding performance in sequence generation on dimeric structures, we compiled the PPIBindBench set, a heterodimer benchmark dataset consisting of recent PDB structures. The PDB entries released between January 1, 2023 and February 2026 and determined by X-ray diffraction or electron microscopy were collected. Entries containing at least two protein entities were retained, leaving 17,029 initial candidates. Heterodimers were identified based on the following criteria. First, we reconstructed each biological assembly and kept only complexes with exactly two protein chains. This step excludes trimeric and larger assemblies. Second, the complexes with identical UniProt IDs were excluded. Third, protein chains longer than 30 amino acids should have at least 15 interface residues, while shorter peptide chains should have at least 5. The interface residues were identified using a heavy-atom distance threshold of 5.0 Å based on the reconstructed biological assembly structures. After the heterodimer and interface-residue filtering, 2,371 complexes remained. Resolution filtering was applied using thresholds of 2.8 Å or better for X-ray structures and 3.5 Å or better for EM structures, reducing the set to 1,996 complexes.

Components of complexes were designated as binder and target chains based on the following criteria. If only one chain was a peptide, it was designated as the binder and the other was designated as the target. Otherwise, the chain with the higher ratio of interface residues to chain length was designated as the binder. If the ratios were equal, the shorter chain was selected. Binder and target sequences were clustered separately at 30% sequence identity using MMseqs2 [35]. Complexes mapping to the same binder–target cluster pair were considered redundant. The resulting non-redundant dataset consisted of 927 heterodimer complexes. To prevent leakage, binders sharing 30% or higher sequence identity over 80% bidirectional coverage with any training-set sequence were further removed. This leaves 582 complexes. Six of them were modelled as unknown residues throughout, so they carry no native sequence and were excluded. The resulting set is named PPIBindBench and consists of 576 complexes, 197 non-peptide and 379 peptide binders. The set was used as the test set throughout this study.

For inverse folding accuracy evaluation, backbone-only PDB files were generated containing only N, CA, C, and O atoms for both binder and target chains. Side-chain atoms, waters, and ligands were removed. Modified amino acids sharing the standard peptide backbone were retained and mapped to their parent residue types using the AlphaFold 3 CCD mapping [21]. Residue numbering was standardized to consecutive integers without insertion codes. The interface residues were re-indexed accordingly. The final set provides backbone structures, ground-truth sequences, and interface residue annotations. Therefore designed sequences can be scored over the full chain or over the interface residues alone.

### 4.3 Model architecture

The architecture outlined in Section 2.1 is specified in full below. The model input is the backbone atomic coordinates (N, C*α*, C, O) at each residue position *i*. Each residue position is associated with an SPSF profile vector **p***_i_* ∈ ℝ^21^. The 21 dimensions correspond to the 20 standard amino acids and an unknown-residue token (X). Alignment gaps are not counted.

The SPSF profiles are constructed through structure homology search using Foldseek [24]. For each input structure, the query chain is extracted and queried against the Foldseek PDB100 database using `foldseek easy-search`. Setting`--alignment-type 0` restricts alignment to 3Di states and excludes amino acid identity information from the retrieval step. The hits are therefore retrieved by shape alone, without sequence-level bias in the search. The search uses an E-value cutoff of 10 and keeps at most 100 hits per query. The Foldseek reference database contains the query structures themselves. All hits at 100% sequence identity to the query are therefore excluded before profile construction in every evaluation reported here. This exclusion is an artefact of evaluating on structures that are in the reference database. A de novo backbone has no counterpart there. The retained hits are accumulated into a count matrix **C** ∈ ℝ*^L×^*^21^, where *L* is the query length. A Foldseek bit score sums over aligned positions, so a longer or closer structural match scores higher. For each structural hit *k* with bit score *b^k^*, the amino acid aligned at query position *i* contributes *b^k^* as follows:

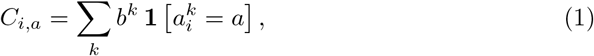

where 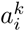 denotes the amino acid aligned to position *i* in hit *k* and **1** is the indicator function, which yields 1 when 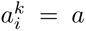, and 0 otherwise. Weighting with bit-score ensures that structurally more similar hits exert greater influence on the profile. Each row is then normalised to yield a per-residue probability distribution **p***_i_* ∈ ℝ^21^:

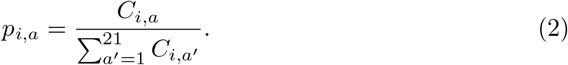

Positions without any aligned residues are set to the zero vector. Profile construction, including the search and the count-matrix assembly, took 4.6 s per structure on eight CPU threads of an AMD EPYC 7542. It was measured over twenty benchmark chains of 33 to 441 residues, 91 s in total, with a per-structure range of 4.0 to 5.2 s. Inference reads the stored profile, so the search is not repeated for each design. On the same twenty chains and one NVIDIA RTX 3090 GPU, ten designs per chain took 95 s in total, excluding profile construction. Including it, the total is 186 s, and the search cost is amortised as more designs are drawn from the same profile.

#### Stage 1: Evolutionary encoder

Following the protein backbone graph convention of [8], the first stage constructs a *k*-NN graph over C*α* atoms with *k*=30. Edge features are computed from 25 pairwise radial basis function (RBF) channels. Each channel encodes the distance between a specific pair of backbone atom types: C*α*–C*α*, N–N, C–C, O–O, C*β*–C*β*, and the remaining 20 directional cross-atom pairs. Each interatomic distance is expanded into 16 Gaussian RBFs. Their centers are evenly spaced from 2 to 22 Å with a width of 1.25 Å. The 25 channels each contribute 16 bins, giving the 400-dimensional RBF descriptor. This descriptor is concatenated with a 16-dimensional relative positional encoding capturing residue offset and chain identity. The signed residue offset *i* − *j* is clipped to ±32 and one-hot encoded into 65 bins. One additional bin marks inter-chain pairs. A linear layer projects this 66-dimensional one-hot vector to 16 dimensions. The resulting 416-dimensional edge input is embedded and lifted to the Stage 1 hidden dimension *d*_s1_ = 256. This assigns the 256-dimensional edge feature vector *e_ij_* to the neighbour pair (*i, j*). The SPSF profile at each node is projected by a linear layer 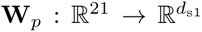 to produce initial node features 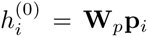. Each of the three encoder layers follows the standard message-passing block with four sub-operations: per-neighbour message computation, message aggregation, a position-wise feed-forward update on the node, and an edge-feature update. For a node *i* with neighbour set *N* (*i*), each layer applies

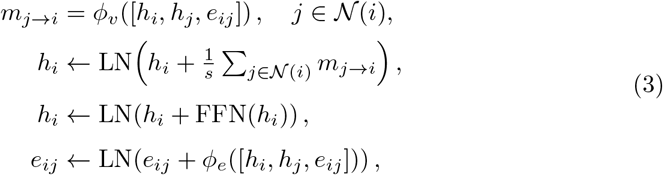

where *ϕ_v_* and *ϕ_e_* are three-layer MLPs, FFN is a position-wise feed-forward network, LN is layer normalisation, and the scaling factor is fixed at *s*=30. All multi-layer perceptrons use GELU activations [36]. The full layer update is provided as Algorithm 1 in Supplementary Methods. The resulting 256-dimensional node features are lifted to *d*_s2_=512 through a linear layer. During training, the initial node features are set to zero for an entire sample with probability *p*_drop_=0.1, so that the model retains structure-only prediction capability.

#### Stage 2: Structure-conditioned MPNN

The second stage builds a denser *k*-NN graph with *k*=48 neighbours and a 512-dimensional edge embedding, computed independently via the same RBF-based edge featurisation. The 512-dimensional node embeddings from Stage 1 serve as initial node features. Three encoder layers refine these features through the same update rules as Stage 1. Sequence decoding proceeds autoregressively through three decoder layers under a per-example random permutation over the positions to be designed [4]. Any fixed-context positions (e.g., target chains in binder design) precede the designed positions in this order. At each decoding step, the node receives messages from two sources: previously decoded neighbours (causal mask, with sequence embeddings), and all other neighbours with their sequence embeddings zeroed (forward mask). The final node representations are projected to 21-dimensional logits and converted to a per-position distribution via softmax (Fig. 1).

Throughout this manuscript, “FoldMapper” refers to this default configuration: *d*_s1_=256. Three alternative variants are evaluated for comparison: (i) a larger evolutionary encoder (*d*_s1_=512), (ii) the BLOSUM62 matrix prior for node feature initialisation, or (iii) both. Foldseek’s own default, alignment-type 2, uses the same matrix during the search. It adds a BLOSUM62 substitution score at temperature *τ* =1.4 to the 3Di structural score [24]. Retrieval here uses alignment-type 0, so that term never enters the search. The BLOSUM variants restore it inside the model instead. The matrix is the same for every query, so it carries no information about the input sequence. What it adds is a prior on plausible substitutions, smoothing each profile position toward biochemically similar amino acids. Concretely, the 20×20 BLOSUM62 matrix [37] is scaled by the same temperature factor *τ* =1.4 and passed through a row-wise softmax. The matrix is reordered to match the Foldseek profile alphabet. The resulting row-stochastic matrix is extended to **B** ∈ ℝ^21^*^×^*^21^ with an identity row/column for the unknown-residue token *X*. Stage 1 node initialisation then takes both the raw profile and its smoothed counterpart:

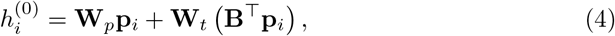

where **B**^⊤^**p***_i_* is the smoothed profile. The matrix **B** is fixed, whereas **W***_p_* and **W***_t_* ∈ ℝ^*d*_*s*1_ × 21^ are trainable.

### 4.4 Training procedure

The default FoldMapper model contains 29.5 M trainable parameters. The model with *d*_s1_=512 contains 40.0 M. All model parameters were initialised with Xavier uniform initialisation [38]. The Adam optimiser [39] was used with *β*_1_=0.9, *β*_2_=0.98, and *ɛ*=10*^−^*^9^. Learning rate followed the Noam schedule [40] with 4,000 warmup steps and a scale factor of 2. The loss for amino acid type prediction was a label-smoothed negative log-likelihood [41] with smoothing weight 0.1. Backbone coordinate augmentation noise was set to *σ*=0.1 Å for the evolutionary encoder and *σ*=0.2 Å for the MPNN encoder. Dropout was set to 0.1 throughout this study. Each mini-batch packed up to 2,000 residues. Each epoch sampled 30,000 training clusters and drew one structure at random from each. Training was performed for 200 epochs on a single NVIDIA RTX 6000 Ada GPU. The default model took about 36 min per epoch, roughly 120 h in total.

### 4.5 Sampling and evaluation

For each structure, we sampled 100 sequences per model under a fixed random seed. The default sampling temperature was set to *T* =0.1, except on antibody–antigen complexes, which used *T* =0.2 to match the source benchmark [42]. By contrast, MapDiff [12] is a discrete diffusion model with no temperature parameter. Its 100 sequences instead arise from independent stochastic DDIM trajectories with Monte-Carlo dropout (Supplementary Methods), so *T* does not apply to it.

Sequence recovery is the fraction of positions at which a sampled amino acid matches the native sequence. In the main text, sequence recovery values are best-of-100 per structure, averaged over structures, except in Section 2.5.

Diversity per structure is the mean pairwise Hamming divergence across the 100 samples:

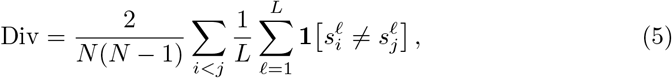

where *N* =100 is the number of samples and *L* is the sequence length. In the main text, diversity is averaged over structures and expressed as percentages.

Perplexity is evaluated on the native sequence *s* via a teacher-forced forward pass:

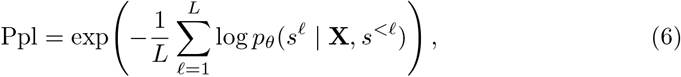

where **X** is the backbone and *s^<ℓ^* the tokens preceding position *ℓ* in the decoding order.

Self-consistency TM-score (scTM) measures whether a designed sequence refolds to the input backbone. Each sampled sequence was folded with AlphaFold 3 [21] and cross-validated with Boltz2 [30]. scTM is the TM-score [43] between the prediction and the reference backbone, computed with USalign [44]. For monomers we used single-chain alignment. For binders, complexes, and oligomers we used multi-chain alignment to obtain a single complex scTM. Model-specific sampling and perplexity protocols are detailed in Supplementary Methods. The scTM normalisation convention is given there as well.

For de novo backbone evaluation, we used 758 backbones from the design set deposited with RFdiffusion [6]. The set comprises 30 monomers, 571 binder complexes, and 157 symmetric oligomers. Every design in that set was experimentally characterised. Therefore, the backbones are expressed design targets rather than arbitrary generated folds. Because these backbones were originally screened with ProteinMPNN-designed sequences, ProteinMPNN may hold a favourable bias on this benchmark. We therefore exclude it from the main comparison (Fig. 3a, b) and report it in Supplementary Table S8.

## Acknowledgements

This work was supported by the Institute of Information & Communications Technology Planning & Evaluation (IITP) grant funded by the Korea government (MSIT) (RS-2023-00220628, Artificial intelligence for prediction of structure-based protein interaction reflecting physicochemical principles). It was also supported by National Research Foundation of Korea (NRF) grants funded by the Korea government (MSIT) (RS-2024-00352229, and RS-2026-25506638 under the Bio&Medical Technology Development Program). Further support came from the Korea Machine Learning Ledger Orchestration for Drug Discovery Project (K-MELLODDY, RS-2026-25608347), funded by the Ministry of Health & Welfare and the Ministry of Science and ICT, Republic of Korea. Support also came from the Korean ARPA-H Project through the Korea Health Industry Development Institute (KHIDI), funded by the Ministry of Health & Welfare, Republic of Korea (RS-2026-25614921). This subject is supported by Korea Ministry of Environment as “The Chemical Accident Prevention Technology Development Project” (RS-2023-00219144). This work was also supported by the Samsung Science & Technology Foundation (C1016338-01-01).

## Author Contributions

H.K. designed the model and conducted the experiments. J.L. supervised the project. All authors discussed the results and contributed to the manuscript.

## Data Availability

The Encyclopedia of Domains (TED) dataset is available from [CATH Encyclopedia of Domains](https://encyclopedia.cathdb.info/) [28]. The ProteinMPNN training and test data are available from the ProteinMPNN GitHub repository (https://github.com/dauparas/ProteinMPNN) [4]. The RFdiffusion experimentally tested backbones used for the foldability evaluation are available at Figshare (DOI: https://doi.org/10.6084/m9.figshare.22662094) [6]. The antibody–antigen structures are available from SAbDab [45] (https://opig.stats.ox.ac.uk/webapps/sabdab-sabpred/sabdab/). The 30 complexes evaluated here follow the benchmark of Li et al. [42]. The two benchmarks constructed in this work are available at Zenodo. TEDMonoBench comprises 10,000 TED domains and 2,126 PDB chains, with the domain identifiers and the redundancy filters applied (DOI: https://doi.org/10.5281/zenodo.24186014). PPIBindBench carries the 576 complexes evaluated here, with the binder and target chain assignments (DOI: https://doi.org/10.5281/zenodo.24186333). The training data is available at Zenodo (DOI: https://doi.org/10.5281/zenodo.24186408). The record holds the cluster splits, the per-chain metadata, and the data-preparation pipeline.

## Code Availability

The FoldMapper source code is available at https://github.com/haelyn-kim/foldmapper under the Apache-2.0 license. The repository holds the model, the training and sampling scripts, the profile generation, and the four configuration files. The trained weights for the four variants are deposited at Zenodo (DOI: https://doi.org/10.5281/zenodo.24186464).

