## Supplementary Information for "FoldMapper: Protein Inverse Folding Model with Structure-Derived Evolutionary Profiles and a Multiscale Graph Encoder"

### Supplementary Information: FoldMapper

#### Supplementary Methods

The core evaluation metrics (sampling, sequence recovery, diversity, perplexity, and scTM) are defined in Methods. This section provides the model-specific protocols and statistical procedures.

##### Evaluation protocols

***Sampling.***

A fixed random seed (19720422) was used across all models and structures to ensure reproducibility, including the autoregressive decoding order in ProteinMPNN and FoldMapper. Autoregressive models (ProteinMPNN [1], ESM-IF [2], FoldMapper) sampled from the softmax distribution at each decoding step. KWDesign [3] computed log-probabilities in a single forward pass and sampled 100 sequences by drawing each position independently from its softmax. Because the distribution factorises over positions, sample diversity is bounded by the per-position entropy. LM-Design [4] performed 100 independent iterative refinement runs per structure. MapDiff [5] performed 100 independent DDIM passes per structure with the `diverse=True` option and dropout enabled. This differs from the authors' original ensemble consensus protocol, which averages logits from 50 stochastic passes to produce a single sequence. We evaluate the per-sample mode because a single consensus sequence cannot be used to compute diversity. MapDiff is also a single-chain model: its input graph is built from one chain, with no chain-break encoding and no mechanism for conditioning on fixed positions. On PPIBindBench it therefore designed the binder chain in isolation, without the target chain that the other general baselines received as structural context.

***Perplexity.***

Perplexity follows the teacher-forced definition in Methods, with the model-specific variations below. ProteinMPNN uses a randomly sampled decoding order; ESM-IF and FoldMapper decode left-to-right. KWDesign conditions on the backbone only; preceding tokens are not used. For LM-Design [4], perplexity follows the formulation in the original conditional masked language modelling code. A single teacher-forced forward pass is run with all designed positions masked, and the negative log-likelihood of the native sequence at those positions is exponentiated. For MapDiff [5], perplexity is computed as the cross-entropy of the native sequence under the mean of $E$ stochastic DDIM trajectories' logits, matching the original implementation. We use $E$=10 rather

Folded structures were predicted with AlphaFold 3 [6] and Boltz2 [7] under default inference parameters. The complex scTM (cTM) was computed over all chains with USalign multi-chain alignment [9] (`-mm 1`). The antibody TM-score (Ab TM) restricts the alignment to the designed heavy and light chains. DockQ [13] was computed between the predicted complex and the native structure with the `DockQ` package. Heavy and light chains were treated as the receptor and the antigen as the ligand. Mean and best-of-100 values are computed per entry, then averaged across entries.

For the antibody-specific baselines, AbMPNN [14] was run with the released weights and the published inference script. AntiFold [15] was run in the joint heavy and light chain setting that matches our full HL mode. The general inverse-folding baselines (ProteinMPNN, ESM-IF, LM-Design, KWDesign, MapDiff) followed the same protocols as the TEDMonoBench and PPIBindBench evaluations. As a single-chain model, MapDiff could not be conditioned on the antigen or on the partner chain. Heavy and light chains were generated in separate runs, with the design positions of each chain taken from the corresponding chain of the reference structure. The other models conditioned on the full complex, with the antigen held fixed.

#### Profile quality metrics

The following metrics are used to characterise the structural position-specific frequency (SPSF) profiles constructed via Foldseek and to analyse their relationship with model performance.

##### *Bitscore.*

The bitscore is a normalised alignment score reported by Foldseek for each structural hit. It measures the statistical significance of the structural similarity between the query and hit, independent of database size. Higher bitscores indicate closer structural homology. For each query protein, we record the maximum bitscore across all retrieved hits (*max bitscore*) and the mean bitscore (*mean bitscore*). The max bitscore reflects the quality of the single best structural match available to the model.

##### *Profile entropy.*

For each residue position $i$, the Shannon entropy of the SPSF profile is defined as

$$H_i = -\sum_{a=1}^{21} p_{i,a} \ln p_{i,a}, \tag{1}$$

where $p_{i,a}$ is the probability of amino acid $a$ at position $i$, over the 20 standard amino acids and the unknown-residue token $X$. Low entropy indicates a sharp, peaked distribution (strong evolutionary constraint); high entropy indicates a flat, diffuse distribution (weak constraint). The per-protein entropy is the mean of $H_i$ over the positions covered by at least one retrieved hit. Uncovered positions are excluded because they carry no distribution to summarise.

***Length-controlled partial Spearman correlation.***

Because sequence length correlates with both profile quality metrics and model performance, we report partial Spearman correlations controlling for length. For variables $X$ (profile quality metric) and $Y$ (recovery) with confound $Z$ (sequence length), we compute the three pairwise Spearman correlations $\rho_{XY}$, $\rho_{XZ}$, $\rho_{YZ}$ and combine them using the standard algebraic partial-correlation identity:

$$\rho_{XY\cdot Z} \;=\; \frac{\rho_{XY} - \rho_{XZ}\,\rho_{YZ}}{\sqrt{(1-\rho_{XZ}^2)(1-\rho_{YZ}^2)}}. \tag{2}$$

Significance is assessed via the Fisher $z$-transform, $z = \tanh^{-1}(\rho_{XY\cdot Z})\sqrt{N-3}$, with $N$ the number of observations.

#### Stage 1 encoder layer

The encoder layer update of Stage 1 (Algorithm 1) applies four sub-operations per layer: per-neighbour message computation, message aggregation with a fixed scaling factor, a position-wise feed-forward transformation, and an edge-feature update. The radial-basis and positional features described in Methods form a 416-dimensional edge input. A linear layer projects it to a 128-dimensional edge embedding, which a second linear layer lifts to $\mathbb{R}^{256}$ before the encoder layers. Layer normalisation and residual connections are applied at each sub-block, with dropout (rate 0.1) inserted within each residual branch during training. The scaling factor $s$ is set to 30 to stabilise the magnitude of summed messages. $\phi_v$ and $\phi_e$ are three-layer multi-layer perceptrons applied to the concatenation $[h_i, h_j, e_{ij}]$ ($768 \to 256 \to 256 \to 256$), with output dimension $\mathbb{R}^{256}$ matching the node and edge feature sizes used in the encoder. FFN is a two-layer multi-layer perceptron with hidden expansion factor 4 ($256 \to 1024 \to 256$). All multi-layer perceptrons ($\phi_v$, $\phi_e$, and FFN) use GELU activations [16].

**Algorithm 1** Stage 1 encoder layer (single iteration). Input: node features $\{h_i\}$, edge features $\{e_{ij}\}_{j\in\mathcal{N}(i)}$. Output: updated $\{h_i\}, \{e_{ij}\}$.

**for** $i = 1$ to $L$ **do**
    **for** $j \in \mathcal{N}(i)$ **do**
        $m_{j\to i} \leftarrow \phi_v([h_i, h_j, e_{ij}])$ ▷ message
    **end for**
    $h_i \leftarrow \mathrm{LN}(h_i + \frac{1}{s}\sum_{j\in\mathcal{N}(i)} m_{j\to i})$ ▷ aggregate
    $h_i \leftarrow \mathrm{LN}(h_i + \mathrm{FFN}(h_i))$ ▷ transform
    **for** $j \in \mathcal{N}(i)$ **do**
        $e_{ij} \leftarrow \mathrm{LN}(e_{ij} + \phi_e([h_i, h_j, e_{ij}]))$ ▷ edge update
    **end for**
**end for**

#### Stage 2 encoder layer

Stage 2 encoder layers follow the same four sub-operations as Algorithm 1, with identical layer normalisation, residual connections, dropout (rate 0.1), and scaling factor $s = 30$. The configured sizes differ: node and edge features sit at $d_{s2} = 512$, the $k$-NN size is $k = 48$, $\phi_v$ and $\phi_e$ become three-layer MLPs of shape $1536 \to 512 \to 512 \to 512$, and FFN becomes $512 \to 2048 \to 512$. Unlike Stage 1, the radial-basis edge embedding is produced directly at $\mathbb{R}^{512}$, so no lifting linear layer is required. Edge features are updated at every layer (Algorithm 1).

#### Stage 2 decoder layer

Stage 2 adopts the standard autoregressive graph decoder used in the structured transformer family of inverse folding models [1, 17]. At each decoder layer, node features are updated by combining two message streams: a causal stream conditioned on the sequence embeddings of previously decoded neighbours, and a forward stream where those embeddings are zeroed. The streams are gated by a causal mask and its complement, both derived from the per-example decoding order. The block structure (LN, residual, FFN, dropout, scaling factor $s$) follows the Stage 2 encoder layer above, but no edge-feature update is performed.

#### Architectural variants

Alongside the default FoldMapper ($d_{s1}$=256 without BLOSUM62 bias), we trained three alternative variants to quantify the impact of the evolutionary encoder dimension and of an explicit substitution prior: (i) $d_{s1}$=512; (ii) $d_{s1}$=256 with BLOSUM62 bias; (iii) $d_{s1}$=512 with BLOSUM62 bias. All variants preserve the Multiscale Graph Encoder (MGE) structure: three evolutionary encoder layers ($k$=30, hidden dimension $d_{s1}$) and the MPNN encoder–decoder ($k$=48, hidden dimension 512). Only the Stage 1 hidden dimension and the node initialisation are varied. The tables label these models by trainable parameter count: 29.5 M for $d_{s1}$=256 and 40.0 M for $d_{s1}$=512. The BLOSUM62 prior adds only the $\mathbf{W}_t$ projection, which leaves both figures unchanged at this precision. Setting $d_{s1}$=512 matches the MPNN hidden dimension, so the projection between stages reduces to an identity mapping.

For the BLOSUM62-augmented variants, each residue's SPSF profile $\mathbf{p}_i$ is transformed through the substitution probability matrix $\mathbf{B} \in \mathbb{R}^{21\times21}$. $\mathbf{B}$ is constructed by applying a row-wise softmax with temperature factor $\tau$=1.4 to the 20×20 BLOSUM62 matrix (reordered to the Foldseek amino acid ordering), and extended with an identity row/column for the unknown-residue token $X$. The initial node feature is $h_i^{(0)} = \mathbf{W}_p\,\mathbf{p}_i + \mathbf{W}_t(\mathbf{B}^\top \mathbf{p}_i)$, where $\mathbf{W}_p, \mathbf{W}_t : \mathbb{R}^{21} \to \mathbb{R}^{d_{s1}}$ are learned linear layers and $\mathbf{B}$ is a fixed (non-trainable) buffer. The BLOSUM-smoothed term $\mathbf{W}_t(\mathbf{B}^\top \mathbf{p}_i)$ injects an evolutionary substitution prior, softening the per-position distribution toward biochemically plausible substitutes beyond the raw 3Di-derived profile frequencies.

### Supplementary Tables

**Table S1**: **Mean recovery, perplexity, and diversity across benchmarks (mean aggregation).**

| TEDMonoBench | **TED** ($n$=10,000) | | | **PDB** ($n$=2,126) | | |
|---|---|---|---|---|---|---|
| | Rec. | Ppl. | Div. | Rec. | Ppl. | Div. |
| ProteinMPNN | 40.6 | 5.93 | <u>28.9</u> | 45.5 | 5.14 | <u>28.0</u> |
| ESM-IF | 52.4 | 4.07 | 12.6 | 56.4 | 5.61 | 11.5 |
| KWDesign | 53.4 | 4.61 | 5.1 | **63.6** | 19.55 | 3.7 |
| MapDiff ($n$=10,000/2,119) | 35.5 | 13.25 | **54.6** | 36.7 | 14.15 | **52.3** |
| LM-Design ($n$=10,000/2,125) | 47.3 | 5.25 | 16.1 | 61.2 | 5.35 | 11.3 |
| FoldMapper-29.5M | <u>54.4</u> | <u>3.79</u> | 20.1 | <u>61.5</u> | **3.63** | 13.8 |
| FoldMapper-40.0M | **54.6** | **3.78** | 19.1 | 59.3 | 3.79 | 15.4 |
| FoldMapper-29.5M+BLOSUM | 46.3 | 3.80 | 25.2 | 48.7 | 3.72 | 25.4 |
| FoldMapper-40.0M+BLOSUM | 48.3 | 3.80 | 21.4 | 54.5 | <u>3.70</u> | 17.9 |
| PPIBindBench | **Non-peptide** ($n$=197) | | | **Peptide** ($n$=379) | | |
| | Rec. | Ppl. | Div. | Rec. | Ppl. | Div. |
| ProteinMPNN | 40.3 | 6.38 | 26.7 | 39.1 | 8.42 | 20.0 |
| ESM-IF ($n$=141/369) | <u>46.5</u> | <u>5.37</u> | 14.8 | 33.0 | 11.17 | 15.1 |
| KWDesign | **51.8** | **4.76** | 6.6 | **46.1** | **7.81** | 8.0 |
| MapDiff ($n$=191/377) | 26.9 | 26.2 | **60.6** | 14.5 | 71.9 | **74.4** |
| LM-Design ($n$=196/379) | 24.4 | 5.51 | 16.4 | 15.3 | 17.71 | 11.4 |
| FoldMapper-29.5M | 46.3 | 5.73 | 21.2 | <u>42.7</u> | 8.41 | 17.4 |
| FoldMapper-40.0M | 45.2 | 5.83 | 21.1 | 41.5 | 8.36 | 17.2 |
| FoldMapper-29.5M+BLOSUM | 39.5 | 5.61 | <u>28.0</u> | 37.9 | 8.32 | <u>21.9</u> |
| FoldMapper-40.0M+BLOSUM | 42.6 | 5.73 | 24.4 | 39.9 | <u>8.28</u> | 20.9 |

Mean over per-protein values; $T$=0.1, 100 samples per structure. Unlike the main tables, recovery is the mean per-sample value rather than the best-of-100.

FoldMapper rows use SPSF profiles with every hit at 100% sequence identity to the query excluded (Methods).

For PPIBindBench, metrics are computed on the binder chain; non-peptide entries have >30 aa, peptide entries have $\leq$30 aa.

Per-row $n$ is shown when it deviates from the full set, as TED/PDB (10,000/2,126) for TEDMonoBench and non-peptide/peptide (197/379) for PPIBindBench.

Best results in **bold**; second-best <u>underlined</u>. Values equal at the reported precision are marked jointly.

**Table S2**: **Recovery, perplexity, and diversity across benchmarks (median aggregation).**

| TEDMonoBench | **TED** ($n$=10,000) | | | **PDB** ($n$=2,126) | | |
|---|---|---|---|---|---|---|
| | Rec. | Ppl. | Div. | Rec. | Ppl. | Div. |
| ProteinMPNN | 46.5 | 5.76 | <u>28.6</u> | 51.3 | 4.72 | <u>27.0</u> |
| ESM-IF | 56.6 | 3.90 | 11.9 | 60.8 | <u>3.34</u> | 10.0 |
| KWDesign | 55.8 | 4.41 | 4.9 | **67.0** | **2.95** | 3.0 |
| MapDiff ($n$=10,000/2,119) | 43.1 | 12.18 | **54.3** | 43.2 | 11.76 | **50.9** |
| LM-Design ($n$=10,000/2,125) | 52.3 | 5.09 | 13.7 | <u>65.4</u> | 4.30 | 9.8 |
| FoldMapper-29.5M | <u>59.9</u> | <u>3.70</u> | 20.0 | 62.2 | 3.38 | 13.9 |
| FoldMapper-40.0M | **60.0** | **3.69** | 18.9 | 61.6 | 3.51 | 15.5 |
| FoldMapper-29.5M+BLOSUM | 52.1 | 3.71 | 25.1 | 52.8 | 3.47 | 25.0 |
| FoldMapper-40.0M+BLOSUM | 53.6 | 3.71 | 21.3 | 56.9 | 3.44 | 18.6 |
| PPIBindBench | **Non-peptide** ($n$=197) | | | **Peptide** ($n$=379) | | |
| | Rec. | Ppl. | Div. | Rec. | Ppl. | Div. |
| ProteinMPNN | 45.6 | 5.73 | 26.5 | **50.0** | 7.06 | 19.8 |
| ESM-IF ($n$=141/369) | <u>52.4</u> | <u>4.69</u> | 13.5 | <u>41.2</u> | 8.43 | 13.9 |
| KWDesign | **55.7** | **4.19** | 6.3 | **50.0** | **5.78** | 7.1 |
| MapDiff ($n$=191/377) | 33.9 | 20.1 | **60.1** | 37.5 | 40.0 | **76.0** |
| LM-Design ($n$=196/379) | 25.4 | 4.85 | 13.3 | 14.3 | 8.56 | 9.4 |
| FoldMapper-29.5M | 48.8 | 5.36 | 22.7 | **50.0** | 6.72 | 16.9 |
| FoldMapper-40.0M | 49.1 | 5.27 | 21.4 | **50.0** | 6.86 | 15.9 |
| FoldMapper-29.5M+BLOSUM | 44.1 | 5.21 | <u>28.0</u> | **50.0** | <u>6.63</u> | <u>22.2</u> |
| FoldMapper-40.0M+BLOSUM | 47.8 | 5.22 | 24.9 | **50.0** | 6.70 | 20.4 |

Median over per-protein values; recovery is the best-of-100 value, as in the main tables. $T$=0.1, 100 samples per structure.

FoldMapper rows use SPSF profiles with every hit at 100% sequence identity to the query excluded (Methods).

For PPIBindBench, metrics are computed on the binder chain; non-peptide entries have >30 aa, peptide entries have $\leq$30 aa. Peptide binders are $\leq$30 aa, so per-protein recovery takes coarse fractional values and the median saturates at 50.0 for most methods.

Per-row $n$ is shown when it deviates from the full set, as TED/PDB (10,000/2,126) for TEDMonoBench and non-peptide/peptide (197/379) for PPIBindBench.

Best results in **bold**; second-best <u>underlined</u>. Values equal at the reported precision are marked jointly.

**Table S3: Ablation study across benchmarks (mean aggregation).**

| TEDMonoBench | **TED** ($n$=10,000) | | | **PDB** ($n$=2,126) | | |
|---|---|---|---|---|---|---|
| Variant | Rec. | Ppl. | Div. | Rec. | Ppl. | Div. |
| FoldMapper-29.5M | 54.4 | 3.79 | 20.1 | **61.5** | **3.63** | 13.8 |
| FoldMapper-40.0M | **54.6** | **3.78** | 19.1 | 59.3 | 3.79 | 15.4 |
| FoldMapper-29.5M+BLOSUM | 46.3 | 3.80 | **25.2** | 48.7 | 3.72 | **25.4** |
| FoldMapper-40.0M+BLOSUM | 48.3 | 3.80 | 21.4 | 54.5 | 3.70 | 17.9 |
| w/o SPSF profile | 53.4 | 3.88 | 21.5 | 47.8 | 4.90 | 24.0 |
| w/o Stage 1 encoder | 47.8 | 4.60 | 25.1 | 59.1 | 3.77 | 18.9 |
| w/o Stage 1 encoder & profile | 46.7 | 4.76 | 26.0 | 45.5 | 5.16 | 27.1 |
| PPIBindBench | **Non-peptide** ($n$=197) | | | **Peptide** ($n$=379) | | |
| Variant | Rec. | Ppl. | Div. | Rec. | Ppl. | Div. |
| FoldMapper-29.5M | **46.3** | 5.73 | 21.2 | **42.7** | 8.41 | 17.4 |
| FoldMapper-40.0M | 45.2 | 5.83 | 21.1 | 41.5 | 8.36 | 17.2 |
| FoldMapper-29.5M+BLOSUM | 39.5 | **5.61** | **28.0** | 37.9 | 8.32 | **21.9** |
| FoldMapper-40.0M+BLOSUM | 42.6 | 5.73 | 24.4 | 39.9 | **8.28** | 20.9 |
| w/o SPSF profile | 41.6 | 6.24 | 24.3 | 42.0 | 8.67 | 17.4 |
| w/o Stage 1 encoder | 44.9 | 5.85 | 23.9 | 40.6 | 8.53 | 19.5 |
| w/o Stage 1 encoder & profile | 40.0 | 6.46 | 26.8 | 40.3 | 8.97 | 19.0 |

Sequence recovery, perplexity, and diversity for FoldMapper variants and ablations (mean over per-protein values; $T$=0.1, 100 samples per structure). Unlike the main ablation table, recovery is the mean per-sample value rather than the best-of-100.

The FoldMapper and "w/o Stage 1 encoder" rows use SPSF profiles with every 100%-identity hit excluded (Methods); the profile-free rows are unaffected by that filter.

For PPIBindBench, metrics are computed on the binder chain; non-peptide entries have >30 aa, peptide entries have ≤30 aa.

Best per column among the four model variants in **bold**; the ablation rows are not ranked. Values equal at the reported precision are marked jointly.

**Table S4: Profile quality–recovery correlations with and without length control.**

| Profile metric | Spearman $\rho$ | Partial $\rho$ (length-controlled) | $p$ (partial) | $n$ |
|---|---|---|---|---|
| Max Foldseek bit score | +0.281 | +0.185 | $4\times10^{-94}$ | 12,096 |
| Profile entropy | −0.152 | −0.115 | $1\times10^{-36}$ | 12,096 |
| Number of Foldseek hits | +0.034 | +0.001 | 0.96 | 12,126 |

Spearman correlation between profile quality metrics and per-protein sequence recovery (mean over the 100 sampled sequences), pooled over both TEDMonoBench subsets. The partial column controls for protein length via the algebraic partial-correlation identity (Supplementary Methods); $p$ is the Fisher $z$ test on the partial correlation.

Profiles exclude every hit at 100% sequence identity to the query, as everywhere else in this work.

Proteins with no Foldseek hit have neither a bit score nor a profile entropy, so the first two rows cover 12,096 of the 12,126 chains. The hit count row uses the full benchmark, because a hit count of zero is still defined.

**Table S5: Structural homolog strength differs between the two TED-MonoBench subsets and with chain length.**

| Length (aa) | $n$ | Max bit score | Profile entropy | Number of hits |
|---|---|---|---|---|
| *TED — predicted domains* | | | | |
| ≤50 | 110/106 | 76.2 | 1.47 | 51.2 |
| 51–100 | 3,197/3,196 | 118.5 | 1.95 | 80.6 |
| 101–150 | 3,329 | 147.7 | 1.94 | 84.5 |
| 151–250 | 2,823 | 203.8 | 1.92 | 87.5 |
| >250 | 541 | 358.8 | 1.91 | 90.3 |
| All | 10,000/9,995 | 164.9 | 1.93 | 84.1 |
| *PDB — experimental structures* | | | | |
| ≤50 | 23/11 | 91.0 | 1.49 | 23.2 |
| 51–100 | 188/175 | 180.6 | 1.74 | 65.2 |
| 101–150 | 327 | 334.2 | 1.80 | 79.7 |
| 151–250 | 521 | 565.0 | 1.62 | 85.3 |
| >250 | 1,067 | 1,340.3 | 1.47 | 90.6 |
| All | 2,126/2,101 | 888.3 | 1.58 | 84.6 |

Profile quality metrics for the SPSF profiles used throughout this work, with every hit at 100% sequence identity to the query excluded. Max bit score is the highest Foldseek bit score retrieved for a query; profile entropy is defined in Supplementary Methods.

The $n$ column gives the number of chains in the bin and, after the slash, the number retrieving at least one hit. Max bit score and profile entropy are means over the latter; the number of hits is a mean over every chain in the bin. Five TED and 25 PDB queries retrieve no hit at all.

A bit score is additive over aligned positions and therefore correlates with chain length (Spearman $\rho = +0.51$ on TED, $+0.81$ on PDB), so each subset is broken down by length. PDB chains average 280 aa against 139 aa for TED, which accounts for part of the aggregate gap. Within matched length bins, the best PDB hit scores 1.2–3.7× higher.

##### Structure self-consistency on natural proteins (TEDMonoBench and PPIBindBench)

**Table S6: Structure self-consistency on TEDMonoBench (TED and PDB sets).**

| AlphaFold 3 | **TED** ($n$=82) | | | | **PDB** ($n$=19) | | | |
|---|---|---|---|---|---|---|---|---|
| | scTM | Best | Succ | pLDDT | scTM | Best | Succ | pLDDT |
| ProteinMPNN | **0.876** | **0.942** | **0.820** | **0.858** | 0.932 | 0.958 | 0.919 | 0.894 |
| ESM-IF ($n$=81/19) | 0.851 | 0.920 | 0.722 | 0.855 | 0.923 | 0.936 | **0.946** | 0.884 |
| KWDesign | 0.858 | 0.917 | 0.758 | 0.853 | 0.941 | 0.955 | 0.915 | **0.914** |
| LM-Design | 0.830 | 0.906 | 0.697 | 0.834 | 0.932 | 0.953 | 0.918 | 0.902 |
| MapDiff ($n$=82/18) | 0.611 | 0.882 | 0.315 | 0.778 | 0.858 | 0.946 | 0.803 | 0.833 |
| FoldMapper-29.5M | 0.854 | 0.926 | 0.755 | 0.845 | 0.944 | 0.961 | 0.945 | 0.908 |
| FoldMapper-40.0M | 0.870 | 0.931 | 0.795 | 0.856 | 0.945 | 0.961 | 0.939 | 0.911 |
| FoldMapper-29.5M+BLOSUM | 0.861 | 0.934 | 0.757 | 0.850 | 0.944 | **0.962** | 0.938 | 0.911 |
| FoldMapper-40.0M+BLOSUM | 0.864 | 0.933 | 0.778 | 0.857 | **0.948** | **0.962** | 0.938 | 0.907 |

| Boltz2 | **TED** ($n$=82) | | | | **PDB** ($n$=19) | | | |
|---|---|---|---|---|---|---|---|---|
| | scTM | Best | Succ | pLDDT | scTM | Best | Succ | pLDDT |
| ProteinMPNN | **0.872** | **0.949** | **0.809** | **0.893** | 0.914 | 0.940 | 0.864 | 0.927 |
| ESM-IF ($n$=78/19) | 0.860 | 0.936 | 0.736 | 0.882 | 0.903 | 0.936 | **0.906** | 0.894 |
| KWDesign | 0.850 | 0.928 | 0.747 | 0.871 | 0.926 | 0.955 | 0.887 | 0.928 |
| LM-Design | 0.828 | 0.911 | 0.690 | 0.854 | 0.920 | 0.949 | 0.877 | 0.917 |
| MapDiff ($n$=82/18) | 0.627 | 0.872 | 0.307 | 0.789 | 0.848 | 0.927 | 0.788 | 0.842 |
| FoldMapper-29.5M | 0.857 | 0.937 | 0.737 | 0.872 | **0.930** | 0.955 | 0.902 | 0.929 |
| FoldMapper-40.0M | 0.867 | 0.943 | 0.760 | 0.880 | 0.905 | 0.955 | 0.888 | 0.927 |
| FoldMapper-29.5M+BLOSUM | 0.861 | 0.940 | 0.745 | 0.878 | 0.923 | **0.960** | 0.884 | **0.935** |
| FoldMapper-40.0M+BLOSUM | 0.863 | 0.940 | 0.759 | 0.878 | 0.919 | 0.956 | 0.884 | 0.927 |

scTM is the mean over 100 samples and Best is the best of 100 per structure. Succ and pLDDT are defined in Supplementary Methods. Self-consistency was evaluated on a 101-structure subsample of TEDMonoBench.

Per-row $n$ as TED/PDB is shown when it deviates from the sampled set, 82/19. Differences reflect entries a model could not process.

Best per column in **bold**; second-best underlined; markers denote numerical rank rather than statistical significance. A marker on a row with a reduced $n$ therefore ranks a value obtained on a smaller entry set than the other rows. Values equal at the reported precision are marked jointly.

**Table S7**: **Structure self-consistency on PPIBindBench.**

| AlphaFold 3 | **Non-peptide** ($n$=59) | | | | | **Peptide** ($n$=41) | | | | |
|---|---|---|---|---|---|---|---|---|---|---|
| | scTM | Best | Succ | $PAE_{int}$ | pLDDT | scTM | Best | Succ | $PAE_{int}$ | pLDDT |
| ProteinMPNN | 0.841 | 0.915 | 0.690 | 12.65 | 0.852 | **0.932** | 0.947 | **0.950** | 8.79 | 0.909 |
| ESM-IF ($n$=47/40) | 0.816 | 0.892 | 0.611 | **12.06** | **0.855** | 0.927 | 0.941 | 0.945 | 9.80 | 0.909 |
| KWDesign | 0.810 | 0.874 | 0.603 | 14.64 | 0.842 | 0.930 | 0.941 | **0.950** | 10.03 | 0.903 |
| LM-Design | 0.813 | 0.887 | 0.597 | 14.33 | 0.841 | 0.925 | 0.937 | 0.940 | 10.86 | 0.899 |
| MapDiff ($n$=56/40) | 0.704 | 0.841 | 0.340 | 20.46 | 0.763 | 0.909 | 0.942 | 0.915 | 12.70 | 0.885 |
| FoldMapper-29.5M | 0.836 | 0.895 | 0.657 | 12.91 | 0.847 | 0.929 | 0.947 | 0.931 | 9.14 | 0.905 |
| FoldMapper-40.0M | **0.850** | 0.910 | 0.714 | 12.69 | 0.851 | 0.929 | 0.945 | 0.932 | 8.81 | 0.906 |
| FoldMapper-29.5M+BLOSUM | 0.849 | **0.918** | **0.724** | 13.15 | 0.847 | 0.929 | **0.948** | 0.946 | 8.96 | 0.908 |
| FoldMapper-40.0M+BLOSUM | 0.840 | **0.918** | 0.698 | 12.89 | 0.847 | **0.932** | 0.946 | 0.946 | **8.74** | **0.912** |
| Boltz2 | **Non-peptide** ($n$=53) | | | | | **Peptide** ($n$=41) | | | | |
| | scTM | Best | Succ | $PAE_{int}$ | pLDDT | scTM | Best | Succ | $PAE_{int}$ | pLDDT |
| ProteinMPNN | **0.827** | **0.915** | **0.673** | 9.65 | 0.900 | 0.937 | **0.965** | 0.951 | 5.94 | 0.945 |
| ESM-IF ($n$=47/40) | 0.798 | 0.886 | 0.573 | 10.98 | 0.902 | 0.937 | 0.962 | **0.960** | 6.00 | 0.940 |
| KWDesign ($n$=55/41) | 0.811 | 0.886 | 0.642 | 11.02 | 0.897 | 0.933 | 0.958 | 0.950 | 6.13 | 0.938 |
| LM-Design | 0.800 | 0.878 | 0.605 | 11.60 | 0.894 | 0.932 | 0.957 | 0.958 | 6.01 | 0.938 |
| MapDiff ($n$=50/40) | 0.724 | 0.850 | 0.384 | 13.67 | 0.838 | 0.920 | 0.958 | 0.946 | 5.87 | 0.937 |
| FoldMapper-29.5M | 0.815 | 0.894 | 0.615 | 10.21 | 0.903 | 0.935 | **0.965** | 0.951 | 6.02 | 0.944 |
| FoldMapper-40.0M | 0.821 | 0.912 | 0.661 | **9.19** | 0.903 | 0.935 | 0.962 | 0.952 | **5.49** | 0.944 |
| FoldMapper-29.5M+BLOSUM | **0.827** | 0.904 | 0.668 | 9.72 | 0.903 | 0.933 | 0.961 | 0.953 | 5.90 | 0.942 |
| FoldMapper-40.0M+BLOSUM | 0.818 | 0.898 | 0.644 | 9.98 | **0.905** | **0.939** | 0.964 | 0.954 | 5.92 | **0.946** |

scTM is the mean complex scTM over 100 samples and Best is the best of 100 per complex. Succ, $PAE_{int}$ and pLDDT are defined in Supplementary Methods. Self-consistency was evaluated on a 100-complex subsample of PPIBindBench.

Non-peptide entries have >30 aa, peptide entries have ≤30 aa.

Per-row $n$ as non-peptide/peptide is shown when it deviates from the predictor's typical set, 59/41 for AlphaFold 3 and 53/41 for Boltz2. Differences reflect entries a model could not process. For ESM-IF these are the twelve longest non-peptide binders, so its non-peptide values come from the shorter subset.

Best per column in **bold**; second-best underlined; markers denote numerical rank rather than statistical significance. A marker on a row whose $n$ differs therefore ranks a value obtained on a different entry set. Values equal at the reported precision are marked jointly.

##### Structure self-consistency on de novo backbones (RFdiffusion, including ProteinMPNN)

**Table S8**: **Structure self-consistency on RFdiffusion de novo backbones (complete results including ProteinMPNN).**

| AlphaFold 3 | **Monomer** ($n$=30) | | | | **Binder** ($n$=571) | | | | **Oligomer** ($n$=157) | | | |
|---|---|---|---|---|---|---|---|---|---|---|---|---|
| | scTM | Best | Succ | pLDDT | scTM | Best | Succ | pLDDT | scTM | Best | Succ | pLDDT |
| ProteinMPNN† | **0.963** | **0.989** | **0.968** | **0.802** | **0.886** | **0.963** | **0.811** | **0.885** | **0.732** | **0.992** | **0.575** | **0.879** |
| ESM-IF | 0.745 | 0.948 | 0.589 | 0.717 | 0.746 | 0.912 | 0.342 | 0.833 | — | — | — | — |
| KWDesign | 0.676 | 0.871 | 0.437 | 0.648 | 0.723 | 0.851 | 0.257 | 0.804 | — | — | — | — |
| LM-Design | 0.594 | 0.810 | 0.312 | 0.615 | 0.735 | 0.881 | 0.309 | 0.814 | — | — | — | — |
| MapDiff | 0.434 | 0.665 | 0.062 | 0.459 | 0.653 | 0.772 | 0.061 | 0.735 | — | — | — | — |
| FoldMapper-29.5M | 0.930 | 0.986 | 0.908 | 0.781 | 0.857 | 0.959 | 0.711 | 0.878 | 0.551 | 0.925 | 0.327 | 0.793 |
| FoldMapper-40.0M | <u>0.944</u> | <u>0.987</u> | <u>0.946</u> | <u>0.782</u> | 0.855 | 0.959 | 0.708 | 0.876 | 0.583 | <u>0.966</u> | 0.366 | 0.821 |
| FoldMapper-29.5M+BLOSUM | 0.765 | 0.951 | 0.621 | 0.767 | <u>0.877</u> | <u>0.962</u> | <u>0.777</u> | <u>0.880</u> | <u>0.605</u> | 0.953 | <u>0.381</u> | <u>0.829</u> |
| FoldMapper-40.0M+BLOSUM | 0.777 | 0.918 | 0.615 | 0.736 | 0.853 | 0.955 | 0.698 | 0.870 | 0.431 | 0.750 | 0.155 | 0.649 |
| Boltz2 | **Monomer** ($n$=30) | | | | **Binder** ($n$=570) | | | | **Oligomer** ($n$=157) | | | |
| | scTM | Best | Succ | pLDDT | scTM | Best | Succ | pLDDT | scTM | Best | Succ | pLDDT |
| ProteinMPNN† | **0.961** | **0.987** | **0.979** | **0.924** | **0.880** | **0.970** | **0.728** | **0.934** | **0.761** | **0.991** | **0.565** | **0.869** |
| ESM-IF | 0.789 | 0.953 | 0.699 | 0.820 | 0.756 | 0.916 | 0.315 | 0.898 | — | — | — | — |
| KWDesign | 0.728 | 0.918 | 0.538 | 0.703 | 0.731 | 0.867 | 0.235 | 0.859 | — | — | — | — |
| LM-Design | 0.648 | 0.876 | 0.379 | 0.717 | 0.743 | 0.899 | 0.292 | 0.869 | — | — | — | — |
| MapDiff | 0.507 | 0.707 | 0.105 | 0.515 | 0.671 | 0.791 | 0.069 | 0.801 | — | — | — | — |
| FoldMapper-29.5M | 0.920 | 0.983 | 0.920 | <u>0.892</u> | 0.845 | <u>0.963</u> | 0.615 | 0.924 | 0.612 | 0.951 | 0.357 | 0.810 |
| FoldMapper-40.0M | <u>0.936</u> | <u>0.984</u> | <u>0.957</u> | 0.889 | 0.843 | 0.961 | 0.607 | 0.924 | 0.651 | <u>0.976</u> | 0.408 | 0.833 |
| FoldMapper-29.5M+BLOSUM | 0.805 | 0.964 | 0.687 | 0.852 | <u>0.869</u> | **0.970** | <u>0.698</u> | <u>0.930</u> | <u>0.658</u> | 0.975 | <u>0.422</u> | <u>0.834</u> |
| FoldMapper-40.0M+BLOSUM | 0.786 | 0.923 | 0.609 | 0.820 | 0.840 | 0.958 | 0.610 | 0.924 | 0.476 | 0.802 | 0.182 | 0.687 |

scTM is the mean over 100 samples and Best is the best of 100 per structure. Succ and pLDDT are defined in Supplementary Methods.

Oligomer results are restricted to models supporting tied sequence design.

Best per column in **bold**; second-best <u>underlined</u>; markers denote numerical rank rather than statistical significance. Values equal at the reported precision are marked jointly.

†ProteinMPNN may have a favourable bias on this benchmark because the RFdiffusion backbones were originally screened with ProteinMPNN-designed sequences.

##### Structure self-consistency on antibody–antigen complexes (subset30)

**Table S9**: **Structure self-consistency on antibody–antigen complexes (subset30, full HL joint design).**

| AlphaFold 3 | cTM | Best cTM | Succ | DockQ | Best DQ | Best Ab TM |
|---|---|---|---|---|---|---|
| ProteinMPNN | <u>0.691</u> | **0.859** | 0.256 | 0.091 | <u>0.376</u> | <u>0.976</u> |
| ESM-IF | 0.689 | 0.801 | 0.240 | 0.095 | 0.271 | 0.969 |
| KWDesign | 0.684 | 0.824 | 0.229 | 0.100 | 0.301 | 0.970 |
| LM-Design | 0.661 | 0.780 | 0.170 | 0.047 | 0.179 | 0.962 |
| MapDiff | 0.653 | 0.807 | 0.173 | 0.057 | 0.277 | 0.964 |
| AbMPNN | **0.696** | 0.843 | <u>0.280</u> | <u>0.138</u> | **0.411** | 0.974 |
| AntiFold | 0.550 | 0.653 | 0.033 | 0.067 | 0.236 | 0.772 |
| FoldMapper-29.5M | 0.685 | 0.821 | 0.265 | 0.119 | 0.364 | 0.952 |
| FoldMapper-40.0M | 0.688 | 0.836 | **0.282** | **0.141** | 0.371 | 0.962 |
| FoldMapper-29.5M+BLOSUM | <u>0.691</u> | 0.842 | 0.253 | 0.119 | 0.351 | **0.977** |
| FoldMapper-40.0M+BLOSUM | 0.680 | <u>0.852</u> | 0.214 | 0.110 | 0.368 | 0.975 |

| Boltz2 | cTM | Best cTM | Succ | DockQ | Best DQ | Best Ab TM |
|---|---|---|---|---|---|---|
| ProteinMPNN | 0.698 | **0.867** | <u>0.242</u> | 0.080 | 0.352 | <u>0.975</u> |
| ESM-IF | 0.700 | 0.838 | 0.230 | 0.098 | 0.300 | 0.970 |
| KWDesign | 0.699 | 0.818 | 0.232 | 0.090 | 0.251 | 0.969 |
| LM-Design | 0.668 | 0.798 | 0.112 | 0.037 | 0.178 | 0.964 |
| MapDiff | 0.675 | 0.825 | 0.153 | 0.050 | 0.267 | 0.967 |
| AbMPNN | **0.713** | <u>0.852</u> | **0.266** | **0.131** | **0.371** | 0.974 |
| AntiFold | 0.567 | 0.683 | 0.012 | 0.065 | 0.233 | 0.769 |
| FoldMapper-29.5M | 0.689 | 0.820 | 0.207 | 0.102 | 0.326 | 0.962 |
| FoldMapper-40.0M | 0.691 | 0.838 | 0.213 | <u>0.109</u> | 0.346 | 0.970 |
| FoldMapper-29.5M+BLOSUM | 0.699 | 0.847 | 0.220 | 0.100 | <u>0.363</u> | **0.976** |
| FoldMapper-40.0M+BLOSUM | <u>0.702</u> | <u>0.852</u> | 0.209 | 0.105 | 0.350 | 0.972 |

cTM is the complex scTM, Ab TM is the antibody TM-score, and DQ abbreviates DockQ [13], all defined in Supplementary Methods. Columns headed Best give the best of 100 samples per complex and the others give the mean.

AbMPNN and AntiFold are antibody-specific baselines; ESM-IF, LM-Design, KWDesign, MapDiff, and ProteinMPNN are general inverse-folding methods. FoldMapper rows use the self-hit-excluded SPSF profiles applied throughout this work (Methods).

Subset30: 30 SAbDab antigen-clustered complexes (Fab and VHH).

Best per column in **bold**; second-best <u>underlined</u>; markers denote numerical rank rather than statistical significance. Values equal at the reported precision are marked jointly.

AntiFold designs the variable fragment only, so its Ab TM is normalised over constant domains it does not design; the low value reflects this coverage difference rather than a difference in structural fidelity.
